# ACE2-Coated Virus-Like Particles Effectively Block SARS-CoV-2 Infection

**DOI:** 10.1101/2023.09.19.558424

**Authors:** Canan Bayraktar, Alisan Kayabolen, Arda Odabas, Ayşegul Durgun, İpek Kok, Kenan Sevinç, Aroon Supramaniam, Adi Idris, Tugba Bagci-Onder

**Author notes:** These authors contributed equally to this work (Canan Bayraktar, Alisan Kayabolen).

## Abstract

A large body of research accumulated over the past three years dedicated to our understanding and fighting COVID-19. Blocking the interaction between SARS-CoV-2 Spike and ACE2 receptor has been considered an effective strategy as anti-SARS-CoV-2 therapeutics. In this study, we developed ACE2-coated virus-like particles (ACE2-VLPs), which can be utilized to prevent viral entry into host cells and efficiently neutralize the virus. These ACE2-VLPs exhibited high neutralization capacity even when applied at low doses, and displayed superior efficacy compared to extracellular vesicles carrying ACE2, in the in vitro pseudoviral assays. ACE2-VLPs were stable under different environmental temperatures, and they were effective in blocking all tested variants of concern in vitro. Finally, ACE2-VLPs displayed marked neutralization capacity against Omicron BA.1 in the Vero E6 cells. Based on their superior efficacy compared to extracellular vesicles, and their demonstrated success against live virus, ACE2-VLPs can be considered as vital candidates for treating SARS-CoV-2. This novel therapeutic approach of VLP coating with receptor particles can serve as proof-of-concept for designing effective neutralization strategies for other viral diseases in the future.

**Graphical Abstract:** In our study, we demonstrate the prevention of SARS-CoV-2 infection through the use of Ace2-coated VLPs.

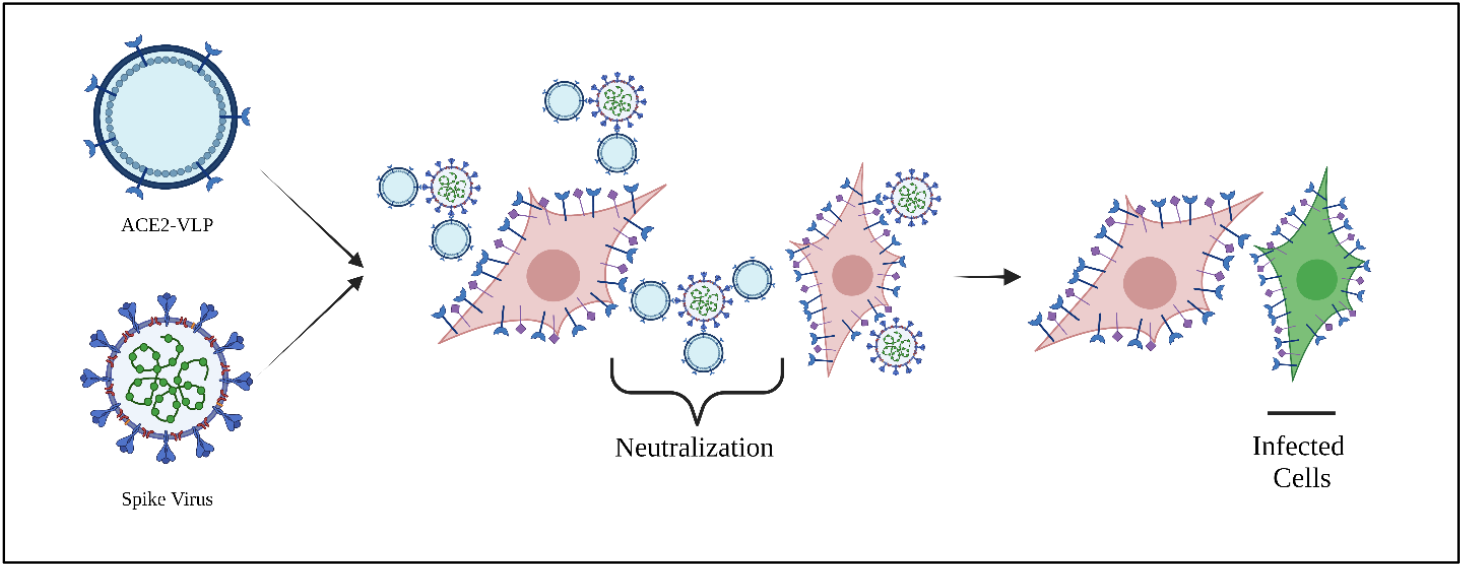

## 1. INTRODUCTION

Molecular strategies that disrupt the direct interaction between microbes and their cognate receptors on host cells may serve as a powerful therapeutic approach for neutralizing a pathogen. Coronavirus disease (COVID-19) is a lethal respiratory disease caused by severe acute respiratory syndrome coronavirus 2 (SARS-CoV-2) [1]. Since the beginning of the global pandemic, extensive research on SARS-CoV-2 biology and therapeutic interventions have revealed novel molecular anti-COVID-19 strategies. SARS-CoV-2 envelope is decorated with the notorious spike protein, which is critical for host cell attachment and entry via host receptor angiotensin-converting enzyme 2 (ACE2) and the priming of the spike protein by the host cell serine protease TMPRSS2 [2]. Hence why blocking the interaction between SARS-CoV-2 spike protein and the ACE2 receptor has been considered an effective antiviral strategy. Current anti-COVID-19 antiviral drugs including molnupiravir and paxlovid are effective at dampening viral replication[3], we have yet to see an approved drug that directly targets the spike protein. Although COVID-19 vaccines have significantly lowered hospitalization and overall mortality [4], the ever changing and evolving COVID-19 variants will pose future challenges, hence warranting better antiviral therapies.

Targeting the interaction between ACE2 and S protein is a plausible antiviral strategy as anti-SARS-CoV-2 therapeutics [5]. Given the upregulated ACE2 levels in COVID-19 patients, the entire ACE2 protein could be a viable target for drug design. Early work by Monteil et al. showed that human recombinant soluble ACE2 (hrsACE2) could significantly block early stages of SARS-CoV-2 infections [6]. Along similar lines, multimeric soluble ACE2 (sACE2) decoy receptors have demonstrated effectiveness in blocking SARS-CoV-2 infection [7-9]. Our recently developed multimerization approach with advanced molecular engineering displayed superior efficacy in ACE2 neutralization in vitro and in vivo, compared to prior multimeric sACE2 based strategies[10]. Despite the potential of sACE2 for SARS-CoV-2 inactivation, high doses of these are needed to achieve the desired effect, which clinically can be problematic for patients [11].

Extracellular vesicles (EVs), lipid bilayered vesicles that are formed by budding from cell membrane, are emerging as an antiviral tool for SARS-CoV-2. Studies have employed EVs decorated with either ACE2 (ACE2-EVs) [5] or anti-SARS-CoV-2 nanobodies [12] to neutralize SARS-CoV-2 via the spike protein. In contrast, virus-like particles (VLPs) offer a more stable, manufacturable, and purifiable method for neutralization. VLPs are non-replicative, non-infective nanostructures consisting of structural viral proteins that mimic native virions but lack viral genetic material, making them an attractive approach for vaccine and therapeutics development[13, 14]. In this study, we developed ACE2-coated virus-like particles (ACE2-VLPs) to prevent SARS-CoV-2 entry into host cells. These ACE2-VLPs were engineered to express more ACE2 on the VLP surface, with the intention to achieve highest neutralization efficiency at low doses. Since host cellular receptors of viruses are not affected by escape mutations[10], we tested ACE2-VLPs on several SARS-CoV-2 pseudo-virus variants. Here, we report that our ACE2-VLPs inhibit the entry of multiple SARS-CoV-2 variants and suggest that ACE2-VLPs have superior neutralizing efficiency compared to ACE2-EVs.

## 2. Materials and Methods

### 2.1. Cell culture

HEK293T cells were purchased from American Type Culture Collection (ATCC, USA) and cultured in DMEM (Gibco, USA), with 10% FBS (Gibco, USA) and 1% Pen/Strep (Gibco, USA) in a 37°C incubator with 5% CO^2^. For transfection experiments, HEK293T cells were seeded on 10-cm plates as 4x10^6^ cells/plate and 6 well plates as 1 x 10^6^ cells/plate. The next day, media were changed, and cells with approximately 90-100% confluency in each plate were transfected with the required plasmid(s) for each assay using 1 mg/ml PEI (polyethyleneimine).

### 2.2. Cloning

ACE2 plasmids with different cytoplasmic tail lengths were generated using the Q5® Site-Directed Mutagenesis (SDM) Kit (New England BioLabs, Ipswich, MA, USA) according to manufacturer instructions. pcDNA3-sACE2(WT)-8his (RRID: Addgene 1492689) plasmid was used as a template, and PCR reaction was performed with specific primers for truncations. PCR products were then incubated with KLD (Kinase, Ligase & DpnI) enzyme mix to circularize them. Plasmids generated via SDM were transformed into E. coli (Stbl3). The resulting plasmids were isolated using a NucleoSpin Plasmid, Mini kit (M&N, Germany).

### 2.3. Virus-like particle (VLP) and extracellular vesicle (EV) production

All transfections were performed with polyethyleneimine (PEI) transfection reagent (Polysciences, 23966-1, Warrington, PA, USA) on HEK293T cells at a 1:4 (w/v) ratio of DNA/transfection reagent. For ACE2-VLP groups, psPAX2 and ACE2-FL; for ACE2-EV groups Fluc-mCherry and ACE2-FL plasmids; for Control-VLP groups Fluc-mCherry and psPAX2 plasmids were prepared at a 1:1 ratio (Fluc-mCherry plasmids were used to equalize the total amount of DNA to be transfected). A total of 7.5 µg of plasmids for 10-cm plates, 2.5 µg for 6 well plates were prepared, added to the transfection reagent mixture, and incubated for 20-30 min at room temperature. Then, the mixture was added to cells dropwise. After 14-16h of transfection, media was removed, cells were washed with PBS, and serum-free DMEM was added to cells. After 48 and 72h of transfection, conditioned media (CM) was collected from plates, and supernatants were filtered with 0.45 µm syringe filters to remove cell debris. CM were either freshly used, kept at 4°C for a maximum of 1-2 days, or concentrated by ultracentrifugation.

### 2.4. Concentrating VLPs and EVs

CMs were collected after 48 and 72h transfection and filtered through 0.45 µm syringe filters. Then samples were loaded into Quick-Seal® round-top polypropylene tubes (Beckman Coulter, CA). Samples were ultracentrifuged at 100,000×g for 70 min at 4°C using the Beckman OptimaTM L-80 XP ultracentrifuge (Beckman Coulter, CA). The supernatant was discarded, and the pellet resuspended in PBS (1:100 of CM) before storing at -80°C.

### 2.5. Size and particle quantification of ACE2-EVs and ACE2-VLPs by the nanoparticle tracking assay (NTA)

ACE2-Evs and VLPs were quantified, and sizes determined using Nanosight 3000 (Malvern Instruments, Worcestershire, UK). All samples were diluted in PBS (1:1000 ratio to a final volume of 1 ml. Three videos for each construct were recorded (60s) and analyzed using NTA 3.4 Build v3.4.003 (Malvern Instruments, Worcestershire, UK), at optimum camera level and different detection thresholds according to sample. The mean size concentration (particles/ml) was calculated.

### 2.6. Characterization of ACE2-EVs and ACE2-VLPs using transmission electron microscopy (TEM)

Carbon coated copper grids were activated by keeping them under UV for 30 minutes in cell culture hood. 4 µL of concentrated sample was dropped onto each grid and dried at room temperature. 5 µL of PBS was added onto grids and left to dry. After washing 3 times, grids were left to dry overnight at room temperature. Images were taken using the Hitachi HT7800 transmission electron microscope (Hitachi, Tokyo, Japan) at 120 kV without any staining.

### 2.7. Immunoblotting

Concentrated EV or VLP total protein concentrations were determined using the Pierce BCA protein assay kit (Thermo Fisher Scientific, USA) These neutralization reagents were mixed with 4X loading dye, which is prepared by mixing 4X Laemmli Sample Buffer (Bio-Rad, USA) with 2-mercaptoethanol in 9:1 ratio and incubated at 95 °C for 10 min. Equal volume of samples were loaded on gradient SDS polyacrylamide gels (Mini-PROTEAN® TGX™ Precast Gels, Bio-Rad, USA) with protein ladder (Precision Plus Protein, Bio-Rad, USA) and run at 30 mA for 75 min. Then, protein transfer was performed viaTrans-Blot® Turbo™ RTA Mini PVDF Transfer Kit (Bio-Rad, USA). The membrane was blocked with PBS-T (0.1% Tween-20) containing 5% nonfat dry milk for 1h with gentle shaking at room temperature. Then, the blocking buffer was replaced with the primary anti-ACE2 antibody (10108-T24, Sino Biological) (1/1000 dilution) and p24-Gag protein (MAB7360-SP, R&D Systems) (2 µg/mL) diluted in PBS-T with 2% BSA and 0.02% NaN3 by gently shaking overnight at 4°C. The next day, the antibody solution was removed, and the membrane was washed 3 times with PBS-T for 5,10, and 15 min. Then, the membrane was incubated with secondary antibody 1:5000 diluted in PBS-T for 1h at RT and washed 3 times with PBS-T for 5,10,15 min. The membrane was incubated with Pierce™ ECL Western Blotting Substrate (Thermo Fisher Scientific, USA) for 2 min at dark and visualized by Odyssey ® Fc Imaging System (LI-COR Biosciences, USA).

### 2.8. SARS-CoV-2 pseudovirus production

Pseudo-viruses were produced as we previously described [12]. HEK293T cells were seeded on 10-cm plates as 4x10^6^ cells/plate. The next day, cells were transfected with Plex-GFP, psPAX2, and Spike-18aa truncated (RRID: Addgene_149541) plasmids. For alpha, beta, gamma, delta, and omicron variants, transfections were made with different spike protein variant plasmids that were produced by site-directed mutagenesis instead of Spike-18aa-WT. After 14-16h of transfection, the medium was removed, and fresh media (DMEM with 10% FBS and 1% Pen/Strep) was added to cells. 48 and 72h after transfection, media was collected, filtered through 0.45µm syringe filters and stored at 4°C for short-term usage.

### 2.9. SARS-CoV-2 pseudovirus neutralization assay

HEK293T cells were transfected with ACE2- and TMPRSS2-expression plasmids (RRID: Addgene_141185 and RRID: Addgene_145843, respectively) to allow infection by pseudo-viruses as previously described [12]. 16h post-transfection, ACE2-TMPRSS2 expressing HEK293T cells were seeded on 96 well plates. The next day, 120 µl of pseudoviruses bearing either WT or different spike protein variants were mixed with different dilutions of either ACE2-EVs or ACE2-VLPs. Mixtures were incubated for 30 min at 37°C before infecting ACE2 and TMPRSS2 expressing HEK293T cells. Infection rate was determined by fluorescence reads on a microplate reader (BioTek’s Synergy H1, VT, USA), as GFP reporter plasmids were packaged during pseudovirus production. Neutralization efficiency was calculated relative to control fluorescence (control-VLP neutralization group). Images were taken by acquiring 2 × 2 images on Cytation5 (BioTek, USA) and analyzed on the Cytation 5 Gen5 software Image Prime v3.10 (Biotek,USA) using manual mode for phase contrast with LED intensity 8, integration time 100 ms, and camera gain 0.5 with a 10× PL FL objective and for GFP 469, 525, LED intensity 6, integration time 22 and camera gain 3 with 10× PL FL objective.

### 2.10. SARS-CoV-2 Omicron BA.1. decoy receptor neutralization assay

The SARS-CoV-2 neutralization assay was performed as previously described [23]. SARS-CoV-2 B.1.1.529 (Omicron) (BA.1sub variant) – VIC35864 initially obtained from the Peter Doherty Institute for Infection and Immunity and Melbourne Health, Victoria, Australia to perform the SARS-CoV-2 neutralization assay and was cultured in Vero E6 cells. ACE2-EVs or ACE2-VLPs were incubated with 250 plaque-forming units (PFUs) of SARS-CoV-2 at the described concentrations for 30 min at room temperature before infecting Vero E6 cells for 1 h at 37°C. A recombinant monoclonal antibody that recognizes SARS-CoV-2 spike protein (CR3022) was used as a neutralizing positive control [37]. The virus was then removed, and the wells layered with 1% methylcellulose viscosity (4,000 centipoises) (Sigma-Aldrich, St. Louis, MO). The numbers of plaques were assessed 4 days after infection at 37°C before fixing in 8% formaldehyde and stained with 1% crystal violet to visualize plaques.

### 2.11. Statistical analysis

All data were analyzed with GraphPad Prism v9 and ImageJ software. Data were presented as mean +/-SD. Two-way ANOVA was used for comparisons, including multiple parameters. A two-sided p-value < 0.05 was considered statistically significant. Details of each analysis are indicated in figure legends.

## 3. Results

### 3.1. Production and optimization of ACE2-VLPs

To produce ACE-2-VLPs, we transfected different viral packaging plasmids and the pcDNA3-FL-WT-ACE2 plasmid (referred to as ACE2-FL-WT) into HEK293T cells and conditioned media (CM) containing ACE2-VLPs collected. Previous studies have shown that the soluble form of ACE2 and ACE2-coated EVs can prevent infection of pseudoviruses bearing the spike protein efficiently [5, 15, 16]. We have taken the same approach in our study by co-incubating ACE2-VLPs with pseudoviruses bearing the ancestral wild type (WT) spike protein using GFP signal intensity as a read out for infection **(Figure 1A)**. As neutralization efficiency can vary with different viral packaging elements, we tested different packaging systems originating from lentiviruses or endogenous retroviruses **(Figure 1B)**, these included using pUMVC [17], psPAX2[18], hPEG10[19] and mPEG10[19]. It is important to note that all the different undiluted ACE2-VLP forms neutralized pseudo-spikes more efficiently than ACE2-EVs and soluble ACE2 **(Figure 1C)**, highlighting the potency of ACE2-VLPs. Upon comparing efficiencies across dilutions, lentiviral packaging using the psPAX2 plasmid was deemed the most efficient **(Figure 1C)**. Therefore, we decided to use ACE2-VLPs produced by psPAX2 for downstream experiments.

**Figure 1.**
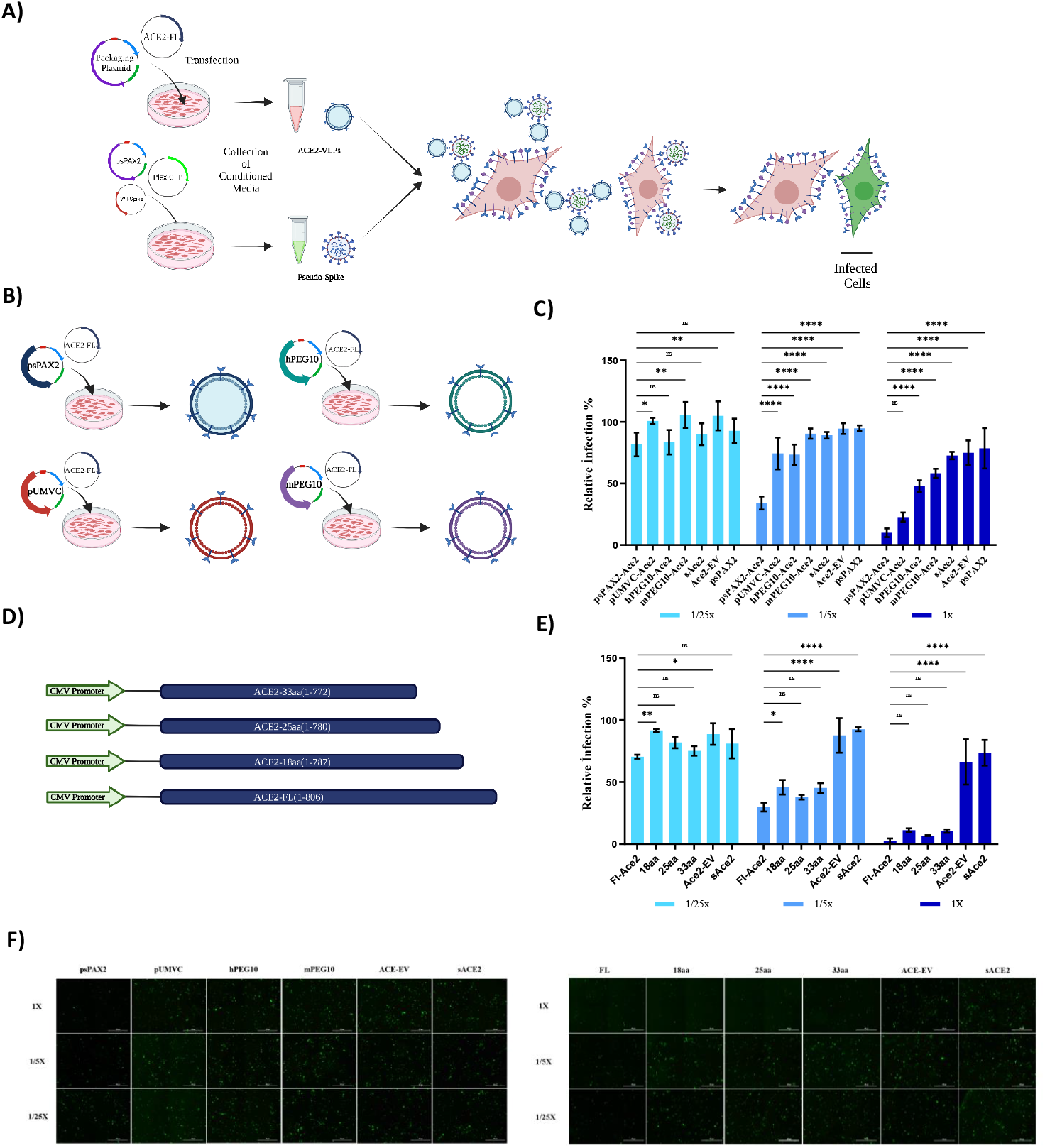
ACE2-VLPs produced using the psPAX2 plasmid lentiviral packaging system bearing full length ACE2 protein had the highest pseudovirus neutralization activity. (A) Schematic representation of ACE2-VLP production and neutralization. (B) Schematic model of the ACE2-VLPs generated by different packaging systems. (Created with Biorender.com) CC) Relative infection rates with pseudovirus in the presence of different dilutions of Conditioned Media (CM) prepared with differently packed ACE2-VLPs. CMs were diluted with DMEM up to 1/25X. CM from cells transfected with psPAX2 alone was used as a control, and infection rates were calculated by measuring relative fluorescence values compared to control wells. (ns: p > 0.05, ^*^: p <= 0.05, ^**^: p <= 0.01, ^***^: p <= 0.001, ^****^: p <= 0.0001, Two way ANOVA.) (D) Schematic representations of ACE2 constructs with different cytoplasmic tail lenghts. (E) Relative infection rates with pseudovirus in the presence of ACE2-coated VLPs with different cytoplasmic tail lengths and dilutions. (F) Representative images of pseudovirus neutralization via ACE2-VLP

We also altered the cytoplasmic tail length of ACE2, to test its effect on the neutralization efficiency. Truncations of 18, 25 or 33 amino acids on the ACE2 cytoplasmic tail were achieved by site-directed mutagenesis **(Figure 1D)** followed by VLP production. ACE2 with the full-length cytoplasmic tail achieved the greatest neutralization efficiency compared to truncated versions as well as soluble ACE2 and ACE2-EV. For example, undiluted full length ACE2-VLP reduced infection down to 2.56 % ± 1.152; while ACE2-EVs could only reduce it down to 29.87 % ± 2.04. Together, these results suggested the functional requirement of the cytoplasmic region of ACE2 for full neutralization activity **(Figure 1E-1F)**.

### 3.2. Characterization and purification of ACE2-VLPs and ACE2-EVs

With the aim of purifying the ACE2-based neutralization agents (EVs and VLPs), which are also needed for clinical application, we concentrated the CM using PEG8000 or single-step ultracentrifugation methods **(Figure 2A)**. Compared to PEG8000 purification, ultracentrifugation yielded higher neutralization efficiency **(Figure S1A)**, and this method proved to be more effective for both ACE2-VLPs and ACE2-EVs than the previously described 4-step ultracentrifugation method [20] [20] **(Figure S1B)**. We then determined the concentration of the ACE2-VLPs and ACE2-EVs **(Figure 2B)**. The median particle diameters were 201.7 nm for ACE2-EV, 172.5 nm for Control-VLPs, and 177 nm for ACE2-VLPs **(Figure 2C)**. We also confirmed the presence of ACE2 in our various constructs by immunoblotting. We observed that there was a marked production of ACE2 protein and no visible difference in the level of packaged ACE2 **(Figure 2D)**. The VLP integrity was also validated by p24-Gag protein, a marker of VLPs, which was only present in Control-VLP and ACE2-VLP samples. Finally, the successful production of ACE2-EVs and ACE2-VLPs was visually confirmed by TEM, where spheres reminiscent of EV and VLP structures were observed **(Figure 2E)**.

**Figure 2.**
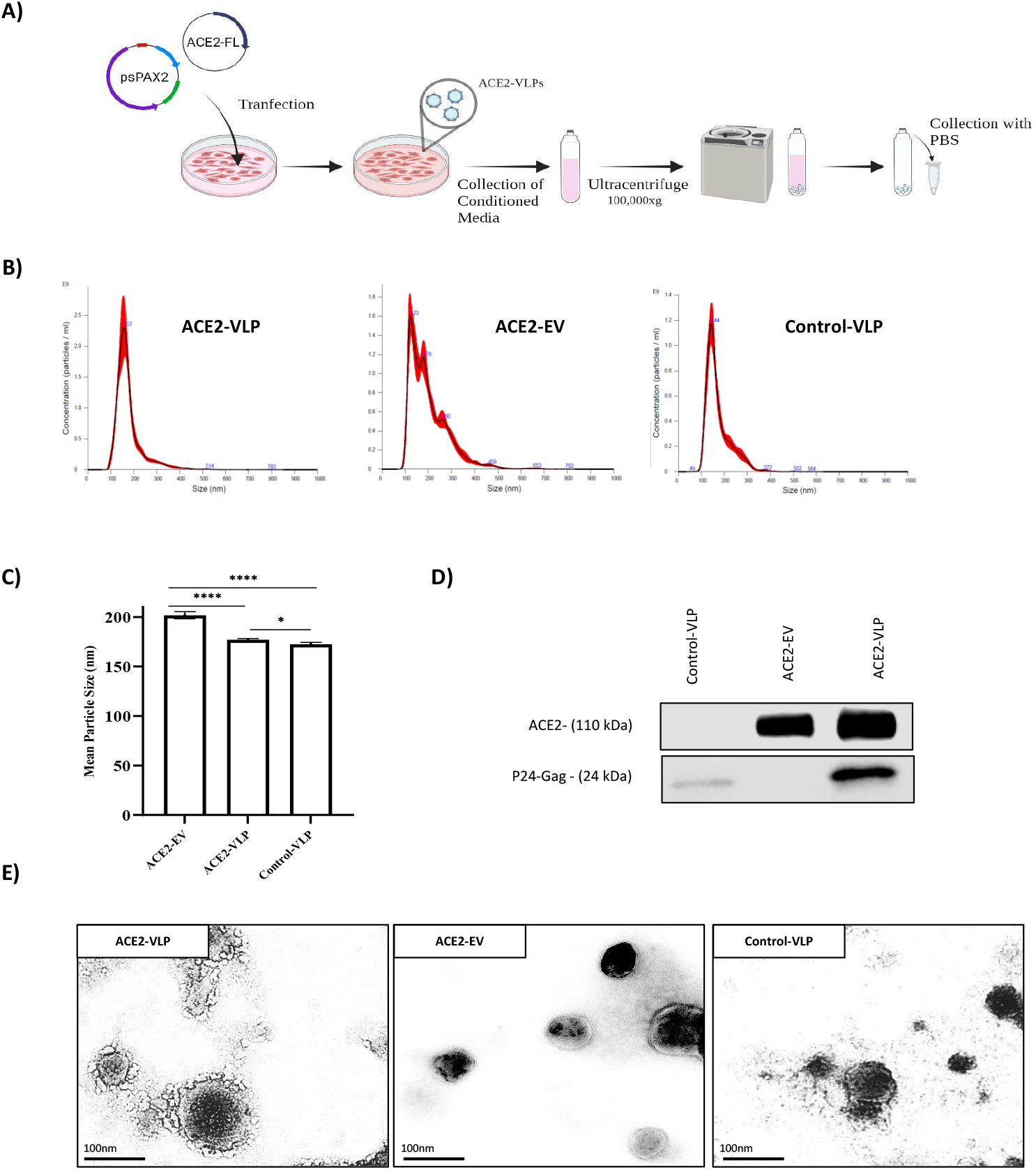
ACE2-VLPs and ACE2-EVs were successfully characterized and purified. (A) Schematic representation of ACE2-VLP production and purification. (B) Size distribution of ACE2-EV, ACE2-VLP and Control-VLP measured by NTA. (C) The median diameters of the ACE2-VLP, ACE2-EV, Control-VLP (D) Western blot images showing ACE2 and capsid p24-Gag protein levels in the ACE2-VLP, ACE2-EV, Control-VLP. (E) SEM images of ACE2-VLP and ACE2-EV. VLPs without envelope (Control-VLP) were used as control. Scale bar: 100 µm. ^*^: p <= 0.05, ^****^: p<= 0.0001, Two way ANOVA.) 18

### 3.3. ACE2-VLPs neutralized SARS-CoV-2 pseudovirus infection more efficiently than ACE2-EVs and are equipotent across all tested SARS-CoV-2 pseudo-variants

To compare the neutralizing capabilities of purified ACE2-EVs and ACE2-VLPs, pseudovirus neutralization experiments were conducted with four different dilutions (1x, 1/3x, 1/9x, 1/27x) of ACE2 preparations **(Figure 3A)**. ACE2-VLPs outperformed ACE2-EVs, providing more efficient neutralization across all tested dilutions **(Figure 3B)**. Notably, even at the highest dilution ACE2-VLPs provided an almost complete pseudovirus neutralization when compared to both ACE2-EVs and control-VLPs **(Figure 3C and 3D)**. Since a potential challenge of VLPs would be to deliver them to patients without sacrificing stability, we tested the stability of the neutralization efficiency of ACE2-VLPs by exposing them to varying environmental temperatures. Remarkably, ACE2-VLPs stored in temperatures as high as 37°C still retained its neutralizing bioactivity **(Figure 3E)**. Together, we showed that ACE2-VLPs are thermo-stable and are potent neutralization agents against SARS-CoV-2 pseudoviruses. To test the broad-range neutralization capacity of ACE2-VLPs, we generated pseudo-viruses using spike plasmids having mutations of six different variant or concern (VOCs) (WT, Alpha, Beta, Gamma, Delta, Omicron), as we have previously done [10] [12]. Irrespective of ACE2-VLP dilution, equipotent neutralizing activity across all tested variants was observed (**Figure 4A, 4B**) further underlining the potency of our engineered ACE2-VLPs. This remarkable neutralization capacity was also evident in fluorescence images (**Figure 4C and S2A**).

**Figure 3.**
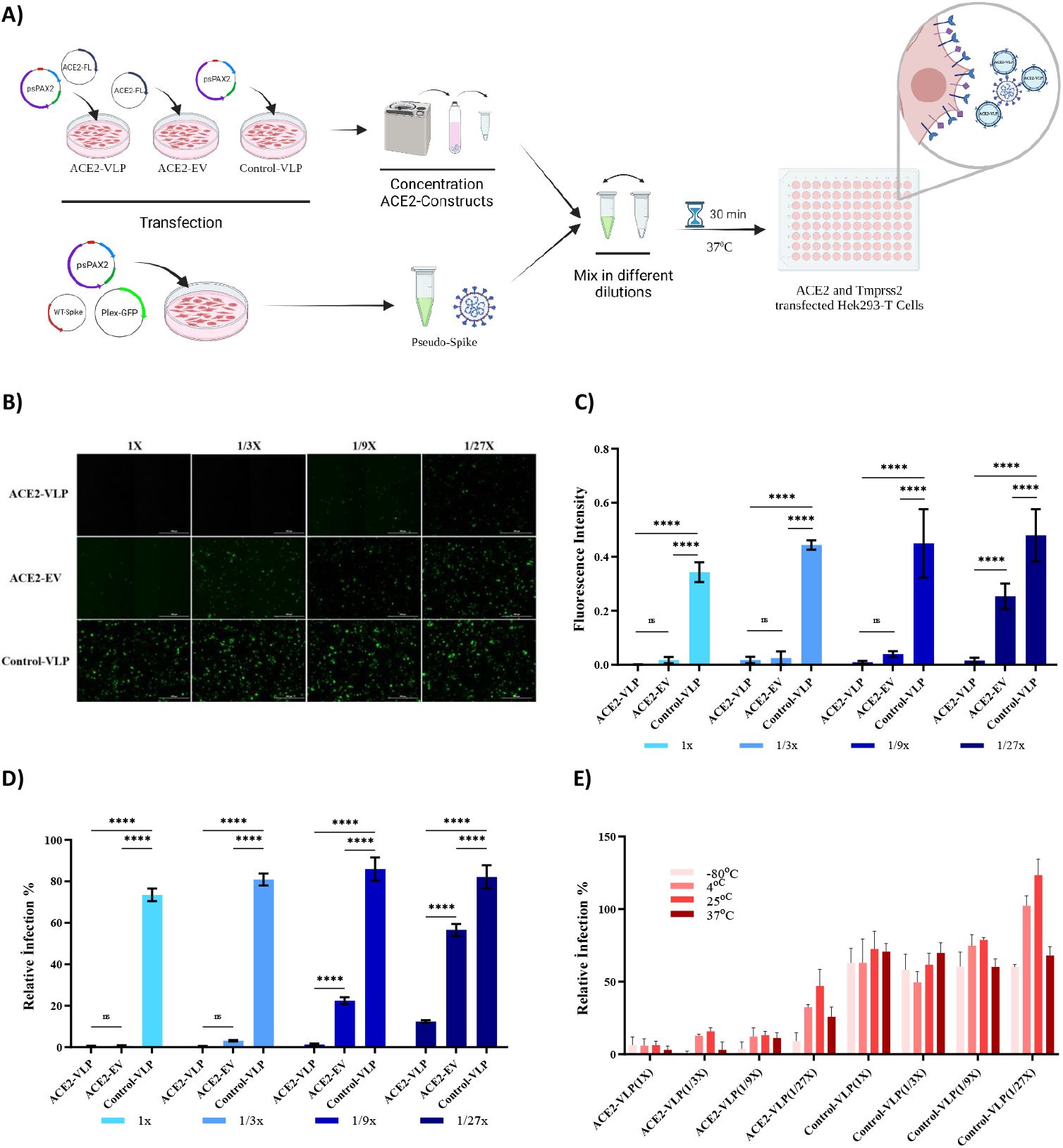
ACE2-VLPs are thermo-stable and are potent neutralization agents against SARS-CoV-2 pseudoviruses. (A) Schematic model of the neutralization experiments with different ACE2 reagents and pseudovirus bearing SARS-CoV-2 spike. (Created with Biorender.com) (B) Representative images of pseudovirus neutralization via ACE2-EV, ACE2-VLP, Control-VLPs. Cells infected with pseudovirus are shown in green. Scale bar: 400 µm (C) Quantification of neutralization. Fluorescence intensities were calculated by analyzing 4 different images for each condition with ImageJ software. (ns: p > 0.05, ^*^: p <= 0.05, ^**^: p <= 0.01, ^***^: p <= 0.001, ^****^: p <= 0.0001, Two way ANOVA) (D) Relative infection rates of pseudoviruses bearing SARS-CoV-2 Spike in ACE2 and TMPRSS2-expressing HEK293T cells in the presence of different dilutions of concentrated ACE2 neutralization reagents. VLPs without envelope (Control-VLP) were used as control. (E) Relative infection rates of pseudoviruses bearing SARS-CoV-2 Spike in ACE2 and TMPRSS2-expressing HEK293T cells in the presence of different dilutions of c1o9ncentrated ACE2-VLPs and different temperatures. VLPs without envelope (Control-VLP) were used as control.

**Figure 4.**
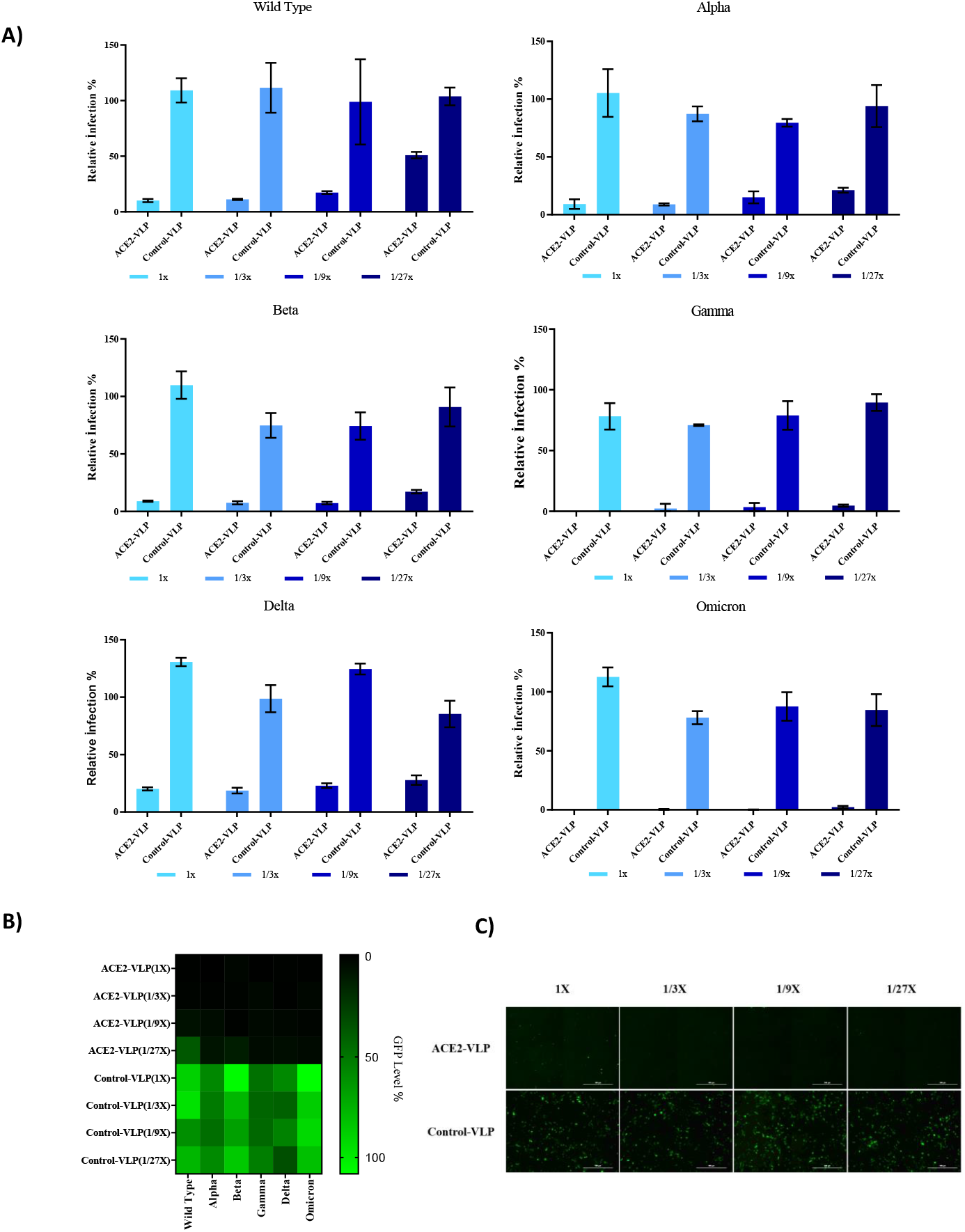
ACE2-VLPs have potent neutralizing activity across all tested SARS-CoV-2 pseudo-variants. (A) Relative infection rates of pseudoviruses bearing VOC-spike in the presence of ACE2-VLP in HEK293T cells expressing ACE2 and TMPRSS2. ACE2-VLPs were serially diluted in culture media. Culture medium was used as control, and infection rates were normalized to fluorescence level of control. (B) Comparison of VOC pseudo-spike neutralization. (C) Representative images of Omicron neutralizations via ACE2 constructs. Scale bar: 400µM

### 3.4. ACE2-VLPs neutralized Omicron BA.1 variant SARS-CoV-2 infections

To measure the neutralization potential of ACE2-VLPs on live virus, we performed a virus neutralization assay using SARS-CoV-2 B.1.1.529 (Omicron), as described previously [12] and measured the viral titer using viral immune plaque assay [21]. When we co-incubated omicron SARS-CoV-2 with ACE2-VLPs, we observed striking neutralization capacity of ACE2-VLPs compared to control-VLP, ACE2-EV and the positive control, a recombinant monoclonal antibody against ancestral spike protein (CR3022). While CR3022 exhibited 49.3 ± 4.1 % neutralization, ACE2-VLPs’s neutralization effect ranged from 100 % to 82.7± 1.5 % from highest (1x) to lowest (1/27x) dilutions. **(Figure 5A-5B)**. Notably, there were no detectable viral plaques in the ACE2-VLP treatment group **(Figure 5C)**. Together, these results showed the potent neutralization capacity of ACE2-VLPs against live SARS-CoV-2 virus.

**Figure 5.**
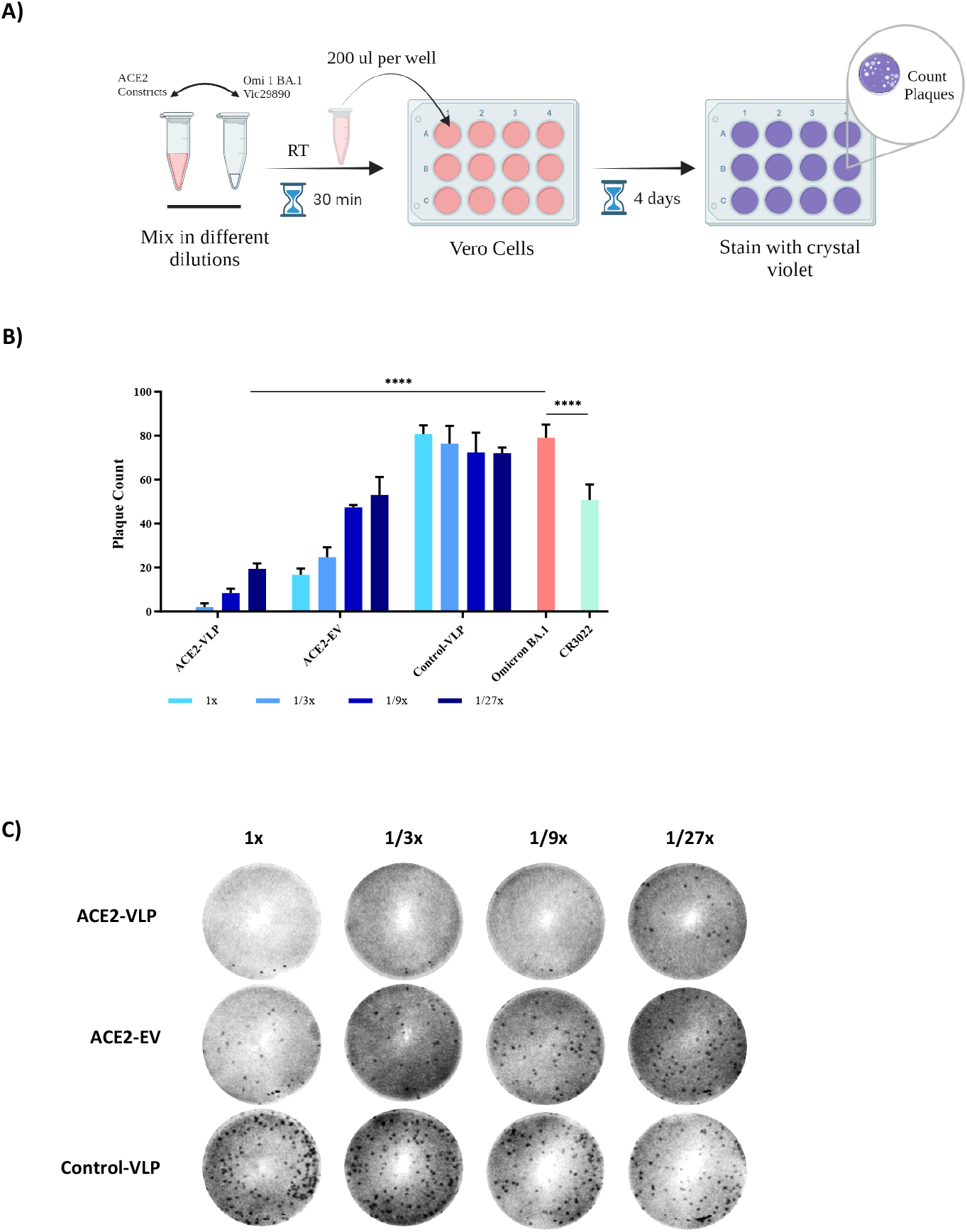
ACE2-VLPs neutralizes live omicron SARS-CoV-2. (A) Schematic model of the experimental setup of Plaque assay. (B) Plaque numbers of Vero-E6 cells with 250 pfu/well Omicron B.A.1 SARS-CoV-2 variant upon 1 h incubation with different dilutions of ACE2 reagents. (C) Representative images for plaques at different dilutions of reagents. (^****^: p <= 0.0001, Two way ANOVA)

## 4. Discussion

As of May 2023, almost 7 million deaths and over 700 million cases were reported about the COVID-19 pandemic. The first infection was reported in Wuhan, China, in December 2019, and COVID-19 was declared a global pandemic by the World Health Organization in March 2020 [22]. Mortality rates have fluctuated based on factors such as age, gender, race, and underlying diseases, and deaths could occur up to 30 days post-infection. More than three years later, the declared global emergency due to the pandemic is over; however, there are still severe cases in need for alternative therapies[1]. The virus is highly contagious and can cause a broad spectrum of symptoms.

To date, there are no specific direct-acting therapies against COVID-19. Furthermore, current antivirals against SARS-CoV-2 are nucleoside analogues and can drive the generation of escape mutants over time[23]. As of March 30, 2023, there were 382 candidate vaccines, 183 of which have been in clinical trials, and as of June 6, 2023, Pfizer-BioNTech, Moderna, and Novavax COVID-19 vaccines were authorized for emergency use and/or FDA-approved [24, 25]. Despite the availability of effective COVID-19 vaccine saving thousands of lives [26], the ability of fast adaptation of SARS-CoV-2 through acquiring new mutations can dampen the effectivity of these vaccines over time. This warrants the development of better direct acting SARS-CoV-2 antivirals.

Previous studies have shown that soluble ACE2 directly binds to SARS-CoV-2 spike protein [16]. Many therapeutic approaches have emerged around ACE2, notably for its ability to bind SARS-CoV-2 and serve as a receptor for its entry into cells [9, 10, 27, 28]. Most of these approaches are limited to the use of ACE2 antibodies. There are disadvantages to this such as harrowing production, low stability, and varying effectiveness against different spike variants. Recombinant protein based ACE2-neutralization approaches have emerged as a potential therapeutic strategy. Though monomeric, dimeric, trimeric, and multimeric recombinant ACE2 decoys have displayed marked neutralization efficacies [10, 29-31], translatability into the clinic remains limited. Due to the high cost involved in the production of producing high purity recombinant proteins [32], this approach has not been the first-line therapy, but the development of these reagent gave us many clues about the biology of virus neutralization.

In this study, we demonstrate that we can successfully prevent SARS-CoV-2 infection *in vitro* using ACE2 protein coated VLPs. VLPs have long been considered useful tools for vaccine development, and VLP-based vaccines have indeed been developed for coronaviruses [33]. Our study ascribes a new use for VLPs as an antiviral agent where VLPs are transformed into ACE2 carriers for direct virus neutralization. A similar study was tested by coating the outer surface of EVs with ACE2 [3, 5]. Given that VLPs are much easier to produce and isolate than EVs, and that they remain more stable and have higher packaging efficiencies than EVs [34], we explored the antiviral capacity of ACE2-coated VLPs. We show that ACE2-VLPs can neutralize SARS-CoV-2 infection more effectively than ACE2-EVs. Importantly, our ACE2-VLPs can effectively neutralize all SARS-CoV-2 pseudovirus VOCs at low concentration, including the live SARS-CoV-2 omicron BA.1 variant.

In conclusion, our approach could be a possible therapeutic strategy for COVID-19 and its future variants. Our work prompts future studies in *in vivo* models, where the ACE2-VLPs that can be stably delivered across different ambient temperatures via various administration routes. Intranasal delivery of such direct neutralizing agents as ACE2-VLPs is an attractive proposal, especially against a virus that is known to replicate within the nasal cavity. Indeed, high SARS-CoV-2 viral load in the nasal cavity have been reported early upon clinical onset[35], a key source of viral aerosol transmission. This will not only pave way for a nasal delivery approach aimed at reducing respiratory COVID-19 illness, but also controlling aerosol SARS-CoV-2 transmission. Our ACE2-VLP strategy will also open opportunities development of effective treatment methods against a broader range of respiratory viruses by multiplexing different envelope proteins against multiple respiratory viruses.

## 5. Conclusion

In summary, our method presents a potential therapeutic solution for both current and upcoming variants of SARS-CoV-2. Our research proves the efficacy of using VLPs coated with ACE2 protein to prevent SARS-CoV-2 infection in vitro. This approach also holds promise for creating treatments against various respiratory viruses by combining different envelope proteins to target multiple pathogens.

## 6. Acknowledgments

We thank Göktu ğ Karabıyık for his help with the image analysis. The authors acknowledge the financial support and use of the services and facilities of the Koç University Research Center for Translational Medicine (KUTTAM). We thank the Molecular Imaging Center and TEM Facility at Koç University N2Star. The CR3022 antibody was obtained from Dr. Naphak Modhiran and Assoc. Prof. Dan Watterson from the School of Chemistry and Molecular Biosciences, The University of Queensland, QLD, Australia. C.B. is supported by a TUBITAK-BIDEB 2211 scholarship for PhD studies.

## 7. Author contributions

Study and experimental design: AK, CB, KS and TBO; cloning: AK and CB; reagent production and concentration; CB,AK, AO, AD, IK; pseudo-virus production and neutralization: AK, CB, AO; western blotting: CB; nanoparticle tracking assay: CB; TEM microscopy: AO; SARS-CoV-2 Omicron BA.1. neutralization: AS, AI; data interpretation: AK, CB, AI and TBO; graphics and figure design: CB; initial manuscript draft: CB, AD, AK and TBO; approved the final manuscript: all authors.

