## Supplemental Figure 1 for "ACE2-Coated Virus-Like Particles Effectively Block SARS-CoV-2 Infection"

### Supporting information

A)

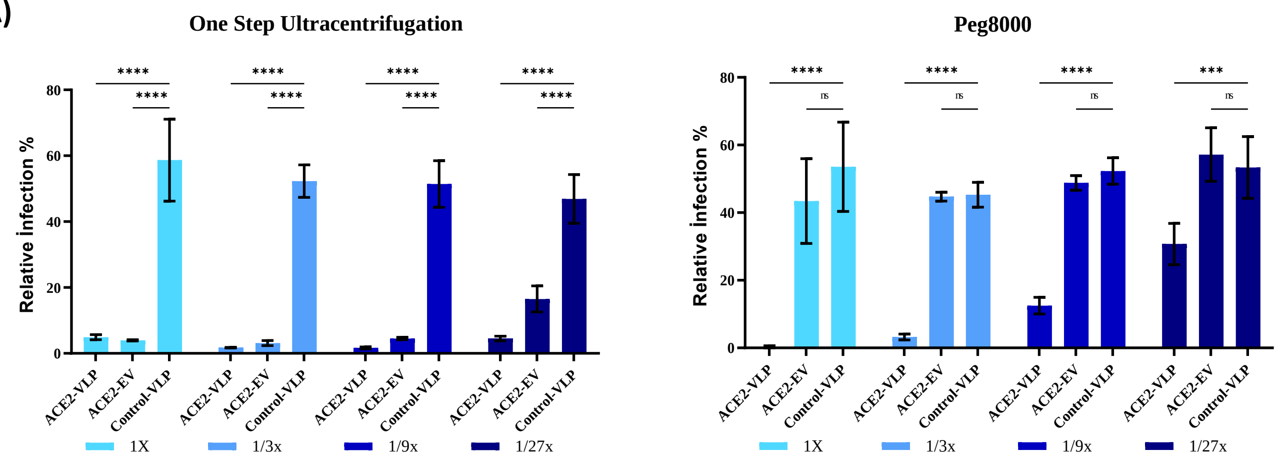


B)

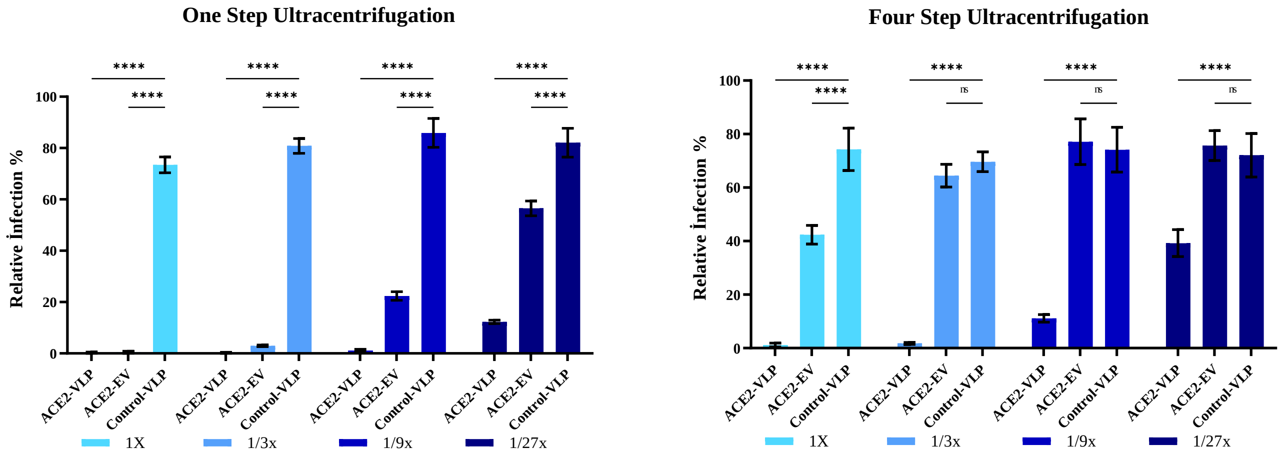


**Supplementary Figure 1. ACE2 reagents purified by single-step ultracentrifugation yielded higher pseudovirus neutralization efficiency.** (A) The relative infection rates of pseudoviruses bearing the SARS-CoV-2 Spike protein were assessed in ACE2 and TMPRSS2-expressing HEK293T cells in the presence of various dilutions of concentrated ACE2 constructs obtained through ultracentrifugation and PEG8000 methods. VLPs without envelope (Control-VLP) were used as control. (B) The relative infection rates of pseudoviruses bearing the SARS-CoV-2 Spike protein were assessed in ACE2 and TMPRSS2-expressing HEK293T cells in the presence of various dilutions of concentrated ACE2 constructs obtained through one-step ultracentrifugation and four-step ultracentrifugation methods. VLPs without envelope (Control-VLP) were used as control. ns: $p > 0.05$, *: $p <= 0.05$, ***: $p <= 0.001$, ****: $p <= 0.0001$, Two way ANOVA.)
